# Sulfo-DIBMA encapsulation uniquely preserves signalling-competent active states of Class B1 GPCRs, CGRP and PTH1 receptors, in native-like nanodiscs

**DOI:** 10.64898/2026.05.13.724797

**Authors:** Farhaan Napier Khwaja, Joseph Gunner, Ellie Thacker, Yasaman Abdolhay, Richard Logan, Philip Kitchen, Dmitry Veprintsev, Mark Wheatley, David Poyner, Hoor Ayub

## Abstract

Class B1 G-protein-coupled receptors (GPCRs), such as the calcitonin gene-related peptide (CGRP) receptor and parathyroid hormone 1 (PTH1) receptor, require native lipid interactions to maintain signalling-competent conformations. However, conventional detergents disrupt these environments. Amphipathic copolymers offer a detergent-free alternative, yet the field still lacks a clear understanding of which polymer architectures best preserve active-state GPCR pharmacology, limiting their broader translational utility. Here, we examine how distinct copolymer chemistries influence the functional integrity of class B1 GPCRs by comparing SMA 2000, DIBMA-12, and the electroneutral sulfo-DIBMA. Using NanoLuciferase bioluminescence resonance energy transfer (NanoBRET) ligand-binding, competition, and mini-G-protein recruitment assays on nanodisc-encapsulated receptors, we show that all three copolymers maintain high-affinity extracellular ligand binding but differ markedly in their ability to preserve intracellular signalling. Despite lower receptor extraction efficiency, only sulfo-DIBMA support mini-Gαs engagement at the CGRP receptor and enable G-protein-dependent allosteric modulation at the PTH1 receptor, including conserved ligand affinity and prolonged residence time. These data reveal that polymer charge and backbone chemistry, rather than extraction yield, determine whether native-like nanodiscs retain the conformational landscape required for active-state signalling. Controlling non-specific ligand binding to the copolymer is a key requirement for a successful assay. Our findings identify sulfo-DIBMALP as a particularly superior environment for preserving native signalling behaviour in class B1 GPCRs, highlighting copolymer chemistry as an important determinant in detergent-free membrane protein studies.

**HIGHLIGHTS:**

- Sulfo-DIBMA encapsulated nanodiscs preserve active-state conformation of human calcitonin gene-related peptide receptor and parathyroid hormone 1 receptor.
- All three copolymers (SMA 2000, DIBMA-12 and sulfo-DIBMA) preserve extracellular ligand binding but only sulfo-DIBMA preserves intracellular functional competence, including mini-Gαs recruitment and G-protein-dependent allosteric modulation.
- Copolymer chemistry, particularly the electroneutral, aliphatic nature of sulfo-DIBMA, may influence the preservation of signalling-competent states in two class B1 GPCRs by minimising charge-driven perturbations during solubilisation.
- Sulfo-DIBMALP provides a novel platform for studying dynamic membrane proteins with potential to provide mechanistic insights and facilitate drug discovery programmes in the future.

**GRAPHICAL ABSTRACT:** 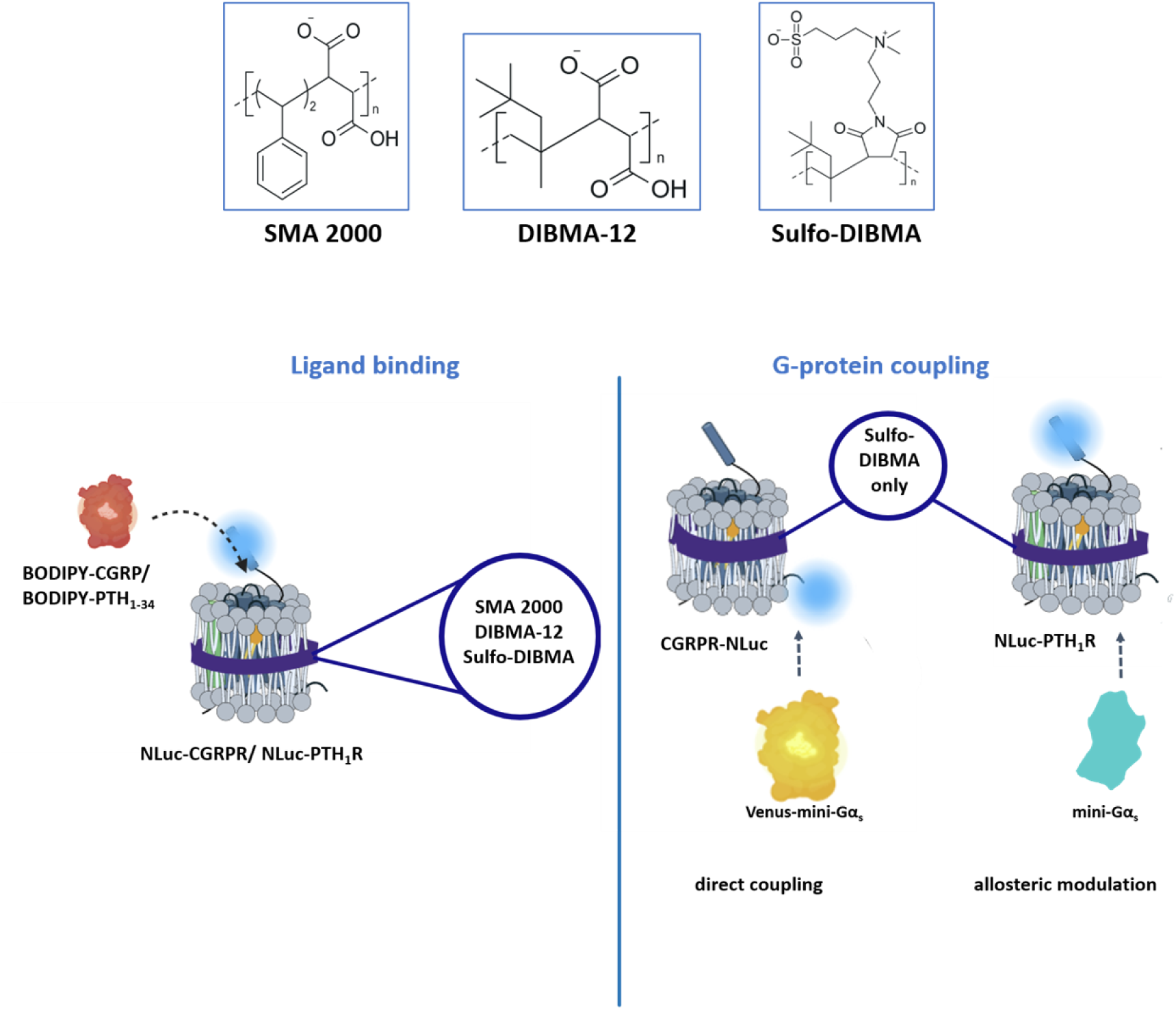

## 1. INTRODUCTION

Integral membrane proteins (IMP) rely on a finely tuned interplay between their transmembrane domains and the surrounding lipid bilayer (1,2). This lipid environment is not a passive scaffold: specific lipid-protein contacts, together with bilayer properties such as thickness, curvature and lateral pressure, actively shape IMP conformation, stability and regulatory behaviour (3–5). Disrupting these interactions destabilises native IMP architecture and compromises function (6), requiring the need for solubilisation strategies that preserve the membrane context.

Detergents remain the most widely used tools for IMP extraction. However, their fundamental limitation is the removal of annular lipids that support native conformational equilibria. By collapsing membrane lateral pressure and stripping away essential lipid species, detergents often shift receptors into non-physiological states, necessitating extensive protein engineering to recover stability (7–9). Amphipathic copolymers offer an attractive alternative (10). Rather than replacing the membrane, they directly extract IMPs together with their endogenous lipid annulus, forming polymer-encapsulated nanodiscs that retain native lipid-interactions (11–13). Because these nanodiscs maintain a physiologically relevant cell membrane environment, they provide a superior platform for accurate structural, biophysical, and pharmacological characterisation of IMPs. This capability will open new opportunities for mechanistic research for drug discovery. To fully realise this potential, it is essential to determine how copolymer architecture governs the extraction of functional IMPs.

Poly(styrene-*co*-maleic acid) (SMA) and poly(diisobutylene-*alt*-maleic acid) (DIBMA) have been used widely for detergent-free solubilisation enabling biophysical studies of class A GPCRs in native-like environments (12,14,15). However, their suitability for class B1 GPCRs remains largely unexplored. This gap is notable given the pharmacological importance of receptors such as glucagon-like peptide-1 receptor, PTH_1_ receptor and CGRP receptor (16–18), and the well-established role of membrane lipids in stabilising peptide ligand binding (as their canonical ligands), extracellular domain orientation and G-protein coupling (18–23). Class B1 GPCRs possess larger extracellular domains, deeper orthosteric ligand binding pockets, and more extensive activation-linked rearrangements particularly around TM6, making them especially sensitive to membrane composition (20,21). Yet it remains unclear how different copolymer chemistries influence the preservation of ligand binding, G-protein engagement and allosteric modulation in these receptors.

To address these gaps, this study evaluates three SMA- and DIBMA-based copolymers as shown in Table 1, namely SMA 2000, DIBMA-12 and sulfobetaine-functionalised poly(diisobutylene-*alt*-maleic acid) (sulfo-DIBMA). This is to capture the potential physiochemical variables likely to influence class B1 GPCR function. SMA 2000, the widely-used aromatic and strongly anionic copolymer (22), provides a benchmark for extraction efficiency and comparison with earlier SMALP studies.

**Table 1.**
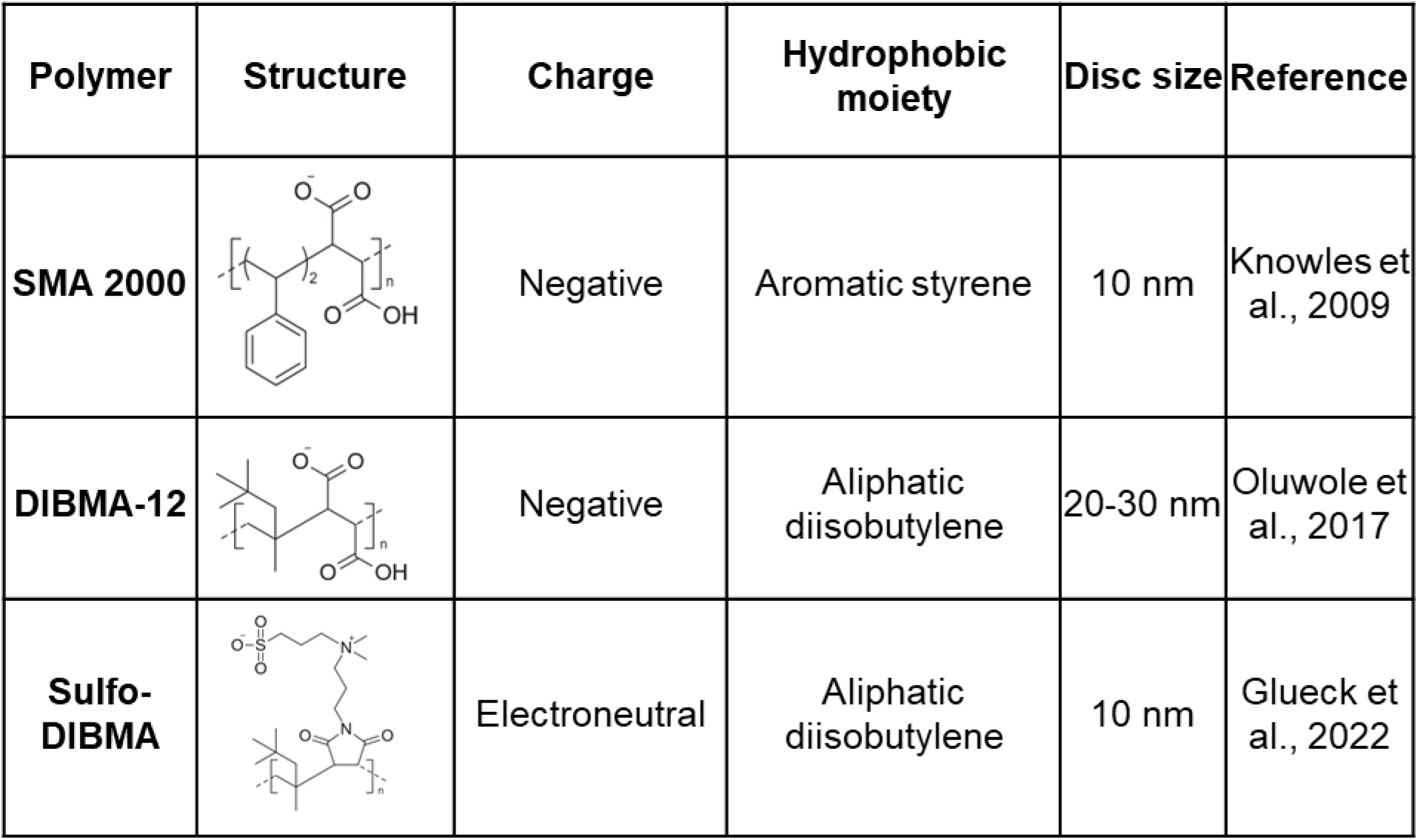
Physicochemical characteristics of the three copolymers used in this study.

| Polymer | Structure | Charge | Hydrophobic moiety | Disc size | Reference |
| --- | --- | --- | --- | --- | --- |
| SMA 2000 |  | Negative | Aromatic styrene | 10 nm | Knowles et al., 2009 |
| DIBMA-12 |  | Negative | Aliphatic diisobutylene | 20-30 nm | Oluwole et al., 2017 |
| Sulfo-DIBMA |  | Electroneutral | Aliphatic diisobutylene | 10 nm | Glueck et al., 2022 |

DIBMA-12, a non-aromatic and less disruptive alternative to SMA (23,24), forms more native-like nanodiscs and enables assessment of whether gentler membrane interactions better preserve receptor conformational flexibility. Sulfo-DIBMA, an electroneutral next-generation copolymer, minimises the charge-related effects of SMA 2000 and DIBMA-12 while retaining the favourable aliphatic DIBMA-like backbone (25), allowing us to test whether reduced electrostatic interference is required for preserving receptor function.

Using these polymers, we examine two therapeutically important class B1 receptors. This included the CGRP receptor, a heterodimer comprising the class B1 GPCR calcitonin receptor-like receptor (CLR) and the accessory protein receptor activity-modifying protein 1 (RAMP1), a validated target in migraine therapy (26,27), and the PTH1 receptor, a prototypical class B1 member and key target in osteoporosis and hypoparathyroidism (28). Together, these receptors provide complementary monomeric and heterodimeric architectures for evaluating copolymer compatibility with class B1 GPCR function.

In this study, NanoBRET-based real-time ligand binding and competition assay (29) revealed that all three copolymers supported high-affinity peptide ligand binding for both CGRP and PTH1 receptors. This demonstrated that copolymer-encapsulated nanodiscs can maintain an intact ligand binding site architecture. However, only sulfo-DIBMA preserved the intracellular conformational rearrangement required for G-protein association and allosteric modulation. This is likely linked to its near-electroneutral chemistry and aliphatic polymer backbone, which may reduce non-specific background and minimise membrane perturbation during solubilisation, thereby supporting a more native-like membrane organisation. Overall, this study identifies sulfo-DIBMA as a particularly compatible platform for examining class B1 GPCR function in native-like lipid environment and offers a mechanistic basis for considering copolymer properties in future pharmacological and structural investigations.

## 2. MATERIALS AND METHODS

### 2.1. Materials

Teriparatide (#4095855, Bachem), PCO371 (#HY-100856, MedChemTronica), DS69910557 (#HY-148350, MedChemTronica), Human α-CGRP (#4013281, Bachem), CGRP 8-37 (#4013696, Bachem), Adrenomedullin (#4034489, Bachem), Adrenomedullin 2 (#4044529, Bachem) olcegepant (#4561, Tocris), rimegepant (#HY-15498, MedChemTronica), and erunumab (#T76874, TargetMol).

Geneticin (#10131027, Thermofisher scientific), DMEM/F12-HEPES (#11330057, Thermofisher scientific), DMEM High glucose w/ L-glutamine w/o sodium pyruvate (Biosera), Polyethylenimine, Branched (#408727 Sigma Aldrich), cOmplete^™^, Mini, EDTA-free protease inhibitor cocktail tablets (#11836170001, Roche), Monoclonal Anti-HA antibody produced in mouse (#H9658-.2ML, Sigma Aldrich), IRDye® 800CW Donkey anti-mouse IgG Secondary antibody (#926-32212, LiCOR). Ni-NTA Agarose (#30210, Qiagen)

Styrene-*co*-maleic acid 1:1 (Cray valley, UK), Styrene-*co*-maleic acid 2:1 (Cray valley, UK), Styrene-*co*-maleimide (Cray valley, UK), Diisobutylene-maleic acid 12, HEPES (#18011, Cube Biotech), and Sulfo-DIBMA (#D-20057, Glycon).

LOBSTR-BL21(DE3)-RIL (#EC1002, Kerafast).

### 2.2. Fluorescent α-CGRP

A custom fluorescent CGRP analogue was constructed by substituting the native Asp^3^ of human α-CGRP with propargylglycine (Pra) to enable site-specific labelling following strategy previously reported for tetramethylrhodamine (TAMRA) labelling (30). The modified peptide was functionalised at Pra with BODIPY 630/650 via alkyne-azide cycloaddition, generating a BODIPY-CGRP ligand. Synthesis was performed using Biosynth custom peptide synthesis service. The peptide sequence and characterisation data are presented in Supplementary Figure 1.

### 2.3. Fluorescent PTH_1-34_

A fluorescent PTH_1-34_ analogue was generated by site-specific labelling at Lys¹³, following the strategy previously reported for TAMRA conjugation (31). The modified peptide was conjugated at the ε-amine of Lys¹³ with BODIPY 630/650, yielding a BODIPY-PTH_1-34_ ligand. Synthesis was performed by Biosynth using their custom peptide synthesis service. The peptide sequence and characterisation data are presented in Supplementary Figure 2.

### 2.4. CGRP receptor cDNA construct design

To enable coordinated expression of CLR and RAMP1, bi-gene CGRP receptor cDNA construct was engineered. A porcine teschovirus-1 2A element (P2A; ATNFSLLKQAGDVEENPGP) self-cleaving peptide sequence (32) was inserted between the human CLR and RAMP1 coding regions to generate a CLR-P2A-RAMP1 construct capable of producing both proteins from one transcript. The design minimises transfection-related variability by ensuring both genes are translated from a single mRNA, thereby promoting tightly coupled expression and simplifying the transfection workflow.

Two custom CGRPR-P2A constructs were synthesised: one containing a Nanoluciferase (NLuc)-tag at the 5’ end of the CLR coding sequence (NLuc-CGRPR) for BRET-based ligand-binding assays, and a second containing the NLuc tag at the 3’ end of the CLR coding sequence (CGRPR-NLuc) for transducer-association assay. Both construct designs and their corresponding amino acid sequences are shown in Supplementary Figure 3.

### 2.5. PTH1 receptor cDNA construct design

A NLuc tagged PTH1 receptor cDNA was generated containing N-terminal His_10_ tag and HA tag followed by the NLuc tag, the human PTH_1_R(_24-593_) coding sequence, Twin-Strep tag sequence at the 3’ end terminating with a stop codon. The cDNA was synthesised by GenScript and cloned into pcDNA 3.1 (+) and pACMV-TetO vector described by (33). PTH_1_ receptor construct designs and corresponding amino acid sequences are shown in Supplementary Figure 4.

### 2.6. Expression constructs

Mini-Gα_s_ and mini-Gα_s/q_ cDNA were cloned into pET15b IPTG inducible vector (34) and Venus-mini-Gα_s_ (35) was cloned into PJ411 vector.

### 2.7. Production of mini-Gα_s_ and mini-Gα_s/q_

Mini-Gα_s_ and Mini-Gα_s/q_ plasmids were expressed and purified according to previously published protocols (36). In brief, a single colony of BL21(DE3)-RIL *E. coli* containing either mini-Gα_s_ or mini-Gα_s/q_ in pET15b was inoculated into 5 ml of Luria–Bertani broth supplemented with 0.2% glucose, 34 μg/ml chloramphenicol and 100 μg/ml ampicillin and incubated at 37 °C, 220 RPM for 4-6 h. This culture was used to inoculate 50 ml of the same media for overnight growth at 30 °C 220 RPM. The next day, the cells were used to inoculate 500 ml of terrific broth supplemented with 0.2 % glucose, 5 mM MgSO_4_, 34 μg/ml chloramphenicol, and 100 μg/ml ampicillin. Cells were allowed to grow at 30 °C, 220 RPM until optical density (OD_600_) reached *ca.* 0.6-0.8. 200 μM of IPTG was used to induce expression and cells were cultured for a further 16-20 h at room temperature (RT), 220 RPM. Cells were then harvested and stored at –80 °C until later use.

The cell pellet was resuspended in buffer (100 mM NaCl, 10 mM imidazole, 10% v/v glycerol, 5 mM MgCl_2_, 5 μM GDP, 100 μM DTT, protease inhibitor tablet, 1 mg/ml lysozyme, 40 mM HEPES; pH 7.5) and stirred at 4 °C for 30 min. The cells were then lysed by sonication (Branson SFX 550) for 10 min with pulses of 20 s on and 40 s off, at 70% amplitude and in an ice bath. The lysate was then pelleted at 35,000 *xg* for 45 min. The supernatant was collected and incubated with Ni-NTA resin for 2-4 h at 4 °C. The resin was then washed with 50 bed volumes of wash buffer (500 mM NaCl, 40 mM imidazole, 10% v/v glycerol, 1 mM MgCl_2_, 5 μM GDP, 20 mM HEPES; pH 7.5) and eluted with high imidazole buffer (100 mM NaCl, 500 mM imidazole, 10% v/v glycerol, 1 mM MgCl_2_, 5 μM GDP, 20 mM HEPES; pH 7.5). The buffer was then exchanged (100 mM NaCl, 1mM MgCl_2_, 5 μM GDP, 10 % v/v glycerol, 10 mM DTT, and 20 mM HEPES; pH 7.5). The sample was then concentrated to around 10 mg/ml, snap frozen, and stored at -80 °C.

### 2.8. Production of Venus-mini-Gα_s_

A single colony of transformed BL21(DE3) *E. coli* containing Venus-mini-Gα_s_ was selected and transfer into 10 ml Luria–Bertani broth supplemented with 50 µg/ml kanamycin and left overnight at 220 RPM, 37°C. The following day, the cells were inoculated into 300 ml of Luria–Bertani broth with 50 µg/ml kanamycin and left to grow at 220 RPM, 37°C until OD_600_ reached 0.6 and then induced with IPTG (1 mM) and incubated over night at lower temperature 220 RPM, 25°C. The next day, induced *E. coli* culture was transferred into 500 ml polycarbonate centrifuge bottles and centrifuged 4500 *xg*, 15 min, supernatant was discarded, and cell pellet was stored in -80 °C for later use or sonicated.

The cell pellet was resuspended in lysis buffer (25 mM Tris/HCl, 150 mM NaCl, 5 mM MgCl₂, 5 µM GDP, protease inhibitor tablet; pH 7.4) and lysed by sonication on ice (Ultrasonic Homogeniser, Qsonica) at 70% amplitude using five cycles of 30 s pulses with 2 min recovery between cycles. The lysate was centrifugated at 4,500 × g for 15 min, and the supernatant was collected and incubated with 2 ml Ni-NTA resin for 4 h at 4 °C. The resin was washed with 50 bed volumes of wash buffer (40 mM imidazole, 25 mM Tris/HCl, 150 mM NaCl, 5 mM MgCl₂, 5 µM GDP; pH 7.4), and bound Venus–mini-Gαs was eluted with elution buffer containing 500 mM imidazole (25 mM Tris/HCl, 150 mM NaCl, 5 mM MgCl₂, 5 µM GDP; pH 7.4). Eluted protein was concentrated using Amicon Ultra centrifugal concentrators (30 kDa MWCO). Full-length fusion proteins were separated from truncation products by size-exclusion chromatography (SEC) on a Superdex 200 10/300 column (Cytiva) using an ÄKTA Pure system. The SEC running buffer comprised 25 mM Tris/HCl, 150 mM NaCl, 5 mM MgCl₂, 5 µM GDP, 5 mM DTT, and 10% glycerol v/v; pH 7.4. Protein concentration was determined by measuring absorbance at 515 nm (Venus excitation maximum) using the extinction coefficient of Venus (104,000 M⁻¹ cm⁻¹) (37).

### 2.9. Cell culture and transfection

HEK 293T cells were maintained in DMEM high glucose supplemented with Fetal Bovine Serum (FBS; 10%). HEK 293T cells were seeded in T175 flasks and grown to 60-80% confluence prior to transfecting with CGRPR-NLuc pcDNA 3.1 (+) using PEI, at a DNA:PEI ratio of 1:3. Cells were cultured for a further 48 h before centrifuging cells at 1000 xg and storing the pellet at –80 °C.

NLuc-PTH_1_R and NLuc-CGRPR in pACMV-TetO mammalian plasmid were transfected into HEK 293S N-acetylglucosaminyltransferase I deficient (GNTI^-^) tetracycline repressor (TetR) cells (HEK 293S GnTI⁻ TetR) using lipofectamine 3000. Stable selection was performed by limiting dilution in the presence of antibiotic geneticin (G418; 2 mg/ml). Monoclonal population of cells were selected, pharmacologically assessed by cAMP accumulation assays and functional populations amplified to express the desired receptor for downstream characterisation.

HEK 293S GnTI⁻ TetR cells stably expressing NLuc-tagged PTH_1_R, wild-type (WT) PTH_1_R, or NLuc-CGRPR were seeded in T175 flasks and grown to 60-80% confluence in supplemented DMEM/F12-HEPES (10% FBS) with 0.5 mg/ml G418. The media was then replaced with supplemented DMEM/F12 without G418 and with tetracycline (2 μg/ml) to induce protein expression by preventing TetR from binding to the tetracycline operator (TetO) site upstream of the coding sequence. Cells were cultured for a further 48-72 h before centrifuging at 1000 xg, discarding the supernatant and storing the pellet at –80 °C.

### 2.10. Preparation of membrane fragments

All steps were performed at 4 °C or on ice. Cell pellets were defrosted and resuspended in ice-cold homogenisation buffer (10 mM Tris-HCl, 250 mM sucrose, and 1 mM EDTA; pH 7.4). Cells were then centrifuged at 1000 xg for 5 min and the supernatant discarded, washed once more with ice-cold homogenisation buffer, centrifuged at 1000 xg for 5 min and the supernatant discarded. The cell pellet was resuspended in homogenisation buffer containing a protease inhibitor tablet. The cells were then lysed on ice by trituration through a 21-guage needle, performing 20-30 passes. The lysate was then centrifuged at 3488 xg for 20 min to removed cellular debris and unbroken cells. The supernatant was collected and then centrifuged at 100,000 xg to collect the membrane fraction. The final pellet was resuspended in buffer A (20 mM HEPES, 50 mM NaCl, 10% glycerol; pH 7) at a membrane concentration of 80 mg/ml and the protein concentration was determined by BCA (4-5 mg/ml). The membrane aliquots were snap frozen and stored at -80 °C until later use.

### 2.11. Solubilisation of membrane fraction

An aliquot of the membrane fraction (80 mg/ml) was allowed to defrost at RT. The membrane fraction was then mixed with a 2x concentrated copolymer stock (10% w/v, 5% w/v, or 2.5% w/v) to give a final copolymer concentration of 5% w/v, 2.5% w/v or 1.25% w/v and a wet membrane concentration of 40 mg/ml. The mixture was then shaken gently for 2-4 h at RT to allow for solubilisation to occur. Following this incubation, the mixture was centrifuged at 100,000 xg at 4 °C for 1 h to separate the insoluble material from the supernatant. The supernatant was then collected, and the insoluble material resuspended in 2% w/v SDS in an equal volume to the supernatant. The supernatant was either snap frozen and stored at –80 °C or incubated on Ni-NTA agarose resin for purification.

### 2.12. Affinity purification

The solubilised fraction was incubated with Ni–NTA agarose resin overnight at 4 °C with gentle agitation, using 500 µl of resin per 1 ml of solubilised material. The mixture was transferred to a gravity-flow purification column, and the flow-through was collected. The resin was washed sequentially with 50 bed volumes of buffer containing 20 mM imidazole, followed by 20 bed volumes of buffer containing 40 mM imidazole, and a final wash with 1 bed volume of buffer containing 60 mM imidazole. Bound receptor was eluted using 3 bed volumes of buffer supplemented with 300 mM imidazole. Elution fractions were pooled and concentrated using a centrifugal concentrator (Vivaspin, 20 kDa MWCO; Sartorius).

### 2.13. BRET-based assays

All measurements were made on the Clariostar plus plate reader (BMG Labtech) equipped with the dual Linear Variable Filter (LVF) monochromator^™^. The emission profile for the NLuc donor was 460 nm-80 and the acceptor was either 660 nm-100 for BODIPY-650/630 or 520 nm-20 for Venus. Each wavelength was read subsequently per well. BRET ratio was defined as the acceptor/donor emission (660/460 or 520/460).

### 2.14. NanoBRET saturation binding assays

Copolymer-encapsulated nanodiscs containing receptors were added to 384-well white optiplates to a final concentration of 0.02-0.2 nM, as calculated from a NLuc standard curve in a final volume of 50 μl of assay binding buffer (150 mM NaCl, 20 mM HEPES, 1 mg/ml BSA, 1 mM EGTA, pH 7.4; or 25 mM HEPES pH 7.4, 104 mM NaCl, 5mM KCl, 1 mM KH_2_PO_4_, 3 mM MgSO_4_, 2 mM CaCl_2_, 1 mg/mL BSA) containing a series of concentrations of BODIPY-PTH_1-34_ or BODIPY-CGRP (in the presence or absence of a saturating concentration of PCO371 or DS69910557 for PTH_1_ receptor or olecegepant for CGRP receptor to determine non-specific binding) for 45 min at 25 °C. Following incubation, 5 μl of assay buffer containing a 1:40 dilution Nano-Glo luciferase substrate (Promega Corporation) was added, followed by an additional 5 min incubation at 25 °C and then reading the BRET measurement. Total binding BRET ratios were corrected for non-specific binding, and the resulting specific binding was fit using a “One-site – specific binding” saturation binding equation in GraphPad Prism 10 to derive dissociation constants (K_d_). Statistical significance was assessed by ordinary one-way ANOVA and Dunnett’s post-hoc analysis against the pooled pK_d_ values, the membrane or buffer pK_d_ was set as the control.

### 2.15. NanoBRET association binding assays

All buffers and plates were pre-chilled prior to running kinetic assays. 2x concentrations of BODIPY-PTH_1-34_ were added to wells of a 384-white optiplate with a buffer only well to baseline correct the data. 2x the concentration of polymer-encapsulated nanodiscs were incubated with mini-Gα protein (3 μM) or equivalent DTT buffer control and with a 1:200 dilution of Nano-Glo luciferase substrate for 10 min prior to loading onto the onboard injectors of the Clariostar Plus plate reader. Association kinetics were initiated once the receptor material was injected into the wells containing the BODIPY-PTH_1-34_, the plate was shaken at 300 RPM for 1 s and the wells read once every 15 s for 10 min. After 10 min, signal decay was apparent, so data was truncated for fitting. Data were fitted with Graphpad Prism 10 to “Association kinetics - Two or more conc. of hot” equation to derive values for K_on_, K_off_, and K_d_ values. Experiments were performed in singlet with at least n=5.

### 2.16. NanoBRET dissociation binding assays

Purified PTH_1_ receptor fractions were incubated with 50 nM (3 μM mini-Gα protein) or 100 nM (buffer control) or BODIPY-PTH_1-34_ for 20 min on ice prior to the addition of 1:200 dilution of Nano-Glo luciferase substrate and incubated for an additional 10 min in a final assay volume of 80 μl. The wells were read once every 15 min for 5 min to capture the baseline prior to the initiation of dissociation by adding 4 μM of unlabelled PTH_1-34_. The plate was then read for another 1 h. The data were fit using Graphpad Prism 10 to a two-phase dissociation model following an Akaike information criteria test. Experiments were performed in duplicates with n=3.

### 2.17. CGRPR-NLuc Venus-mini-Gαs NanoBRET binding assay

Purified CGRP receptor fractions (0.04-0.06 nM) were incubated with 750 nM of Venus-mini-Gαs, a 1:40 dilution of Nano-Glo luciferase substrate and a series of concentrations of α-CGRP, adrenomedullin (AM), and adrenomedullin 2/intermedin (AM2) for 5 min in a final volume of 50 μl prior to reading on the Clariostar Plus plate reader. The plate was read for 30 min in technical singlet with n=3. Data were fit to a “log (agonist) vs. response – Variable slope (four parameters)” in GraphPad Prism 10 and Hillslope, EC_50_ and E_max_ values were derived.

## 3. RESULTS

### 3.1. Comparative solubilisation efficiency of SMA- and DIBMA-like copolymers for CGRP and PTH_1_ receptors

Recombinant human CGRP and PTH₁ receptors stably overexpressed in HEK 293S GnTI⁻ TetR cell membranes were used to assess the solubilisation efficiency of three SMA/DIBMA-based copolymers: SMA 2000, DIBMA-12, and sulfo-DIBMA. An initial screen using 2.5 % w/v SMA 2000 or 5 % w/v of DIBMA-12 or sulfo-DIBMA, revealed differences in their ability to extract each receptor. As shown in Figure 1A and 1C, SMA 2000 efficiently solubilised both receptors, yielding approximately 80% extraction of the CGRP receptor and ∼97% of the PTH₁ receptor. DIBMA-12 showed comparable performance, with ∼80% and ∼98% solubilisation of CGRP and PTH₁ receptors, respectively. In contrast, the electroneutral sulfo-DIBMA copolymer displayed markedly reduced efficiency, extracting only ∼50% of the CGRP receptor and ∼34 % of the PTH₁ receptor.

**Figure 1.**
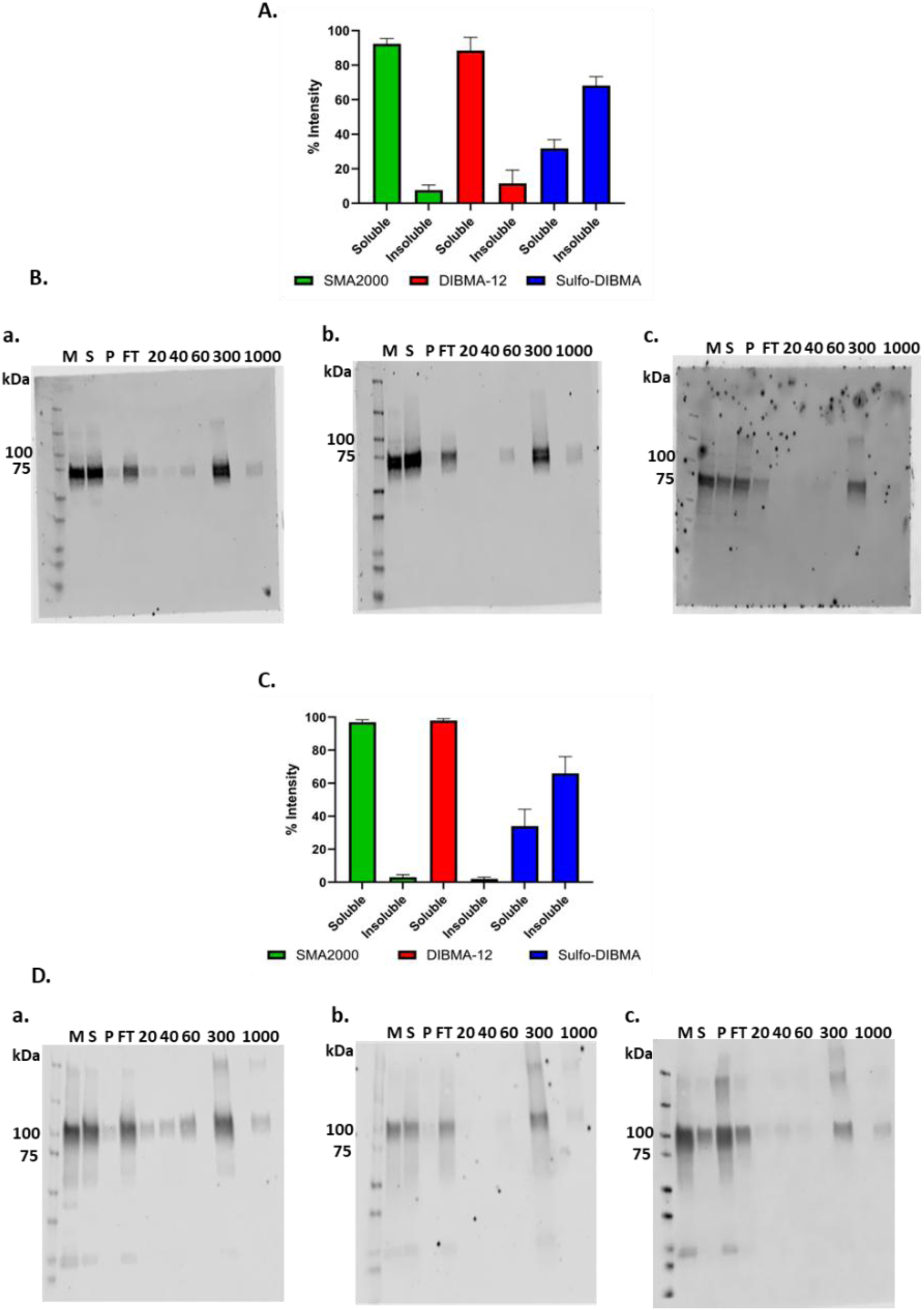
SMA 2000 and DIBMA-12 efficiently solubilise CGRP and PTH₁ receptors, whereas sulfo-DIBMA shows reduced solubilisation efficiency. Recombinant human CGRP and PTH₁ receptors stably overexpressed in HEK 293S GnTI⁻ tetR cell membranes were used to assess the solubilisation efficiency of 2.5% SMA 2000, 5% DIBMA, and 5% sulfo-DIBMA copolymers. Densitometric analysis quantifies receptor solubilisation efficiency using ImageJ, with representative western blots showing solubilised and affinity-purified receptor bands. Anti-HA antibody was used as the primary antibody and IRDye® 800CW Donkey anti-mouse IgG as the secondary. Νluc-PTH1 receptor theoretical size is approximately 92 kDa and Nluc-CLR is approximately 74 kDa. Western blots are representative of n = 3. M= membrane; S= solubilised fraction; P= insoluble pallet; FT= affinity purification flow through. 20, 40, 60, 300 and 1000 show imidazole concentration (mM) wash/elution. (A) Solubilisation efficiency of the three copolymers for the CGRP receptor determined by quantitative densitometric scanning. (B) Representative western blot of solubilised and affinity-purified CGRP receptor using (a) SMA 2000, (b) DIBMA-12, and (c) sulfo-DIBMA. (C) Solubilisation efficiency of the three copolymers for the PTH₁ receptor determined by quantitative densitometric scanning. (D) Representative western blot of solubilised and affinity-purified PTH₁ receptor using (a) SMA 2000, (b) DIBMA-12, and (c) sulfo-DIBMA.

All copolymer-solubilised receptors were subsequently purified by Ni-NTA affinity chromatography via their N-terminal His₁₀-tag. Corresponding western blots and SDS-PAGE gels of solubilised and affinity-purified fractions are shown in Figure 1B and 1D, and in Supplementary Figure 5, respectively. Affinity purification was performed to remove free copolymer and enrich for a ligand-binding-competent receptor population. All three copolymers supported effective purification following solubilisation, and the resulting receptor preparations were subsequently taken forward for functional assessment using NanoBRET-based ligand-binding and G-protein-coupling assays.

### 3.2. Functional retention of CGRP and PTH_1_ receptors in SMALP, DIBMALP and sulfo-DIBMALP

Having established copolymer solubilisation efficiencies, we next assessed whether each nanodisc environment preserved receptor functionality. CGRP and PTH₁ receptors were expressed with an N-terminal NLuc tag (NLuc-CGRPR and NLuc-PTH₁R). To quantify ligand binding-competent conformations, we performed NanoBRET equilibrium saturation binding assays using BODIPY-CGRP (for NLuc-CGRPR) and BODIPY-PTH_1-34_ (for NLuc-PTH₁R) as fluorescent tracers. Non-specific binding was defined in the presence of 10 nM olcegepant for CGRP receptor and 100 µM PCO371 for PTH₁ receptor.

NanoBRET saturation curves in Figure 2 show that all three copolymer nanodiscs (SMALP 2000, DIBMALP and sulfo-DIBMALP) support tracer ligand binding for both receptors. The measured pK_d_ values for BODIPY-PTH_1-34_ at NLuc-PTH₁R and BODIPY-CGRP at NLuc-CGRPR did not differ significantly from values obtained in native membranes, as listed in Table 2, indicating that receptor pharmacology is preserved across nanodisc types.

**Figure 2.**
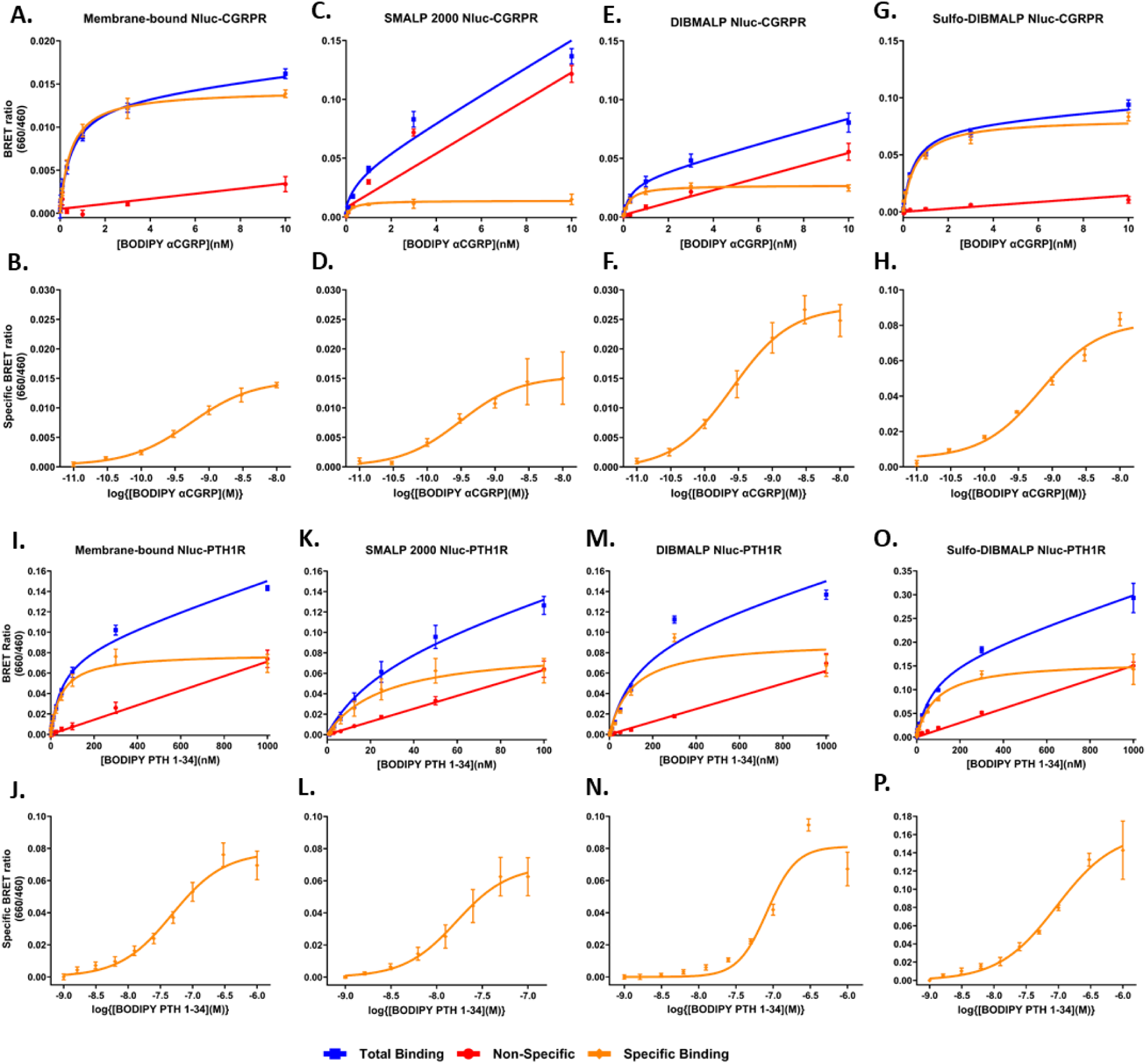
Purified SMALP, DIBMALP, and sulfo-DIBMALP support functional retention of CGRP and PTH₁ receptors. NanoBRET-based equilibrium saturation binding analyses were performed for NLuc-CGRPR (panels A–H) and NLuc-PTH₁R (panels I–P) in membrane preparations and after encapsulation into three copolymer nanodisc environments using BODIPY-CGRP (CGRPR) and BODIPY-PTH1-34 (PTH₁R) as fluorescent tracers. Non-specific binding was defined in the presence of 10 nM olcegepant for CGRPR and 100 µM PCO371 for PTH₁R. For each condition the top panel shows total, non-specific and specific binding fitted to a hyperbolic binding isotherm (panels A, C, E, G for the NLuc-CGRPR and I, K, M, O for the NLuc-PTH1R) and the paired lower panel in each case shows specific binding plotted as a log-sigmoidal binding isotherm (panels B, D, F, H for the NLuc-CGRPR and J, L, N, P for the NLuc-PTH1R). The four columns of panels present data for (left to right); membranes, SMALP 2000, DIBMALP, sulfo-DIBMALP respectively. Data are presented as mean ± s.e.m. from three independent experiments, each performed in technical triplicate (n = 3).

**Table 2.** Polymer-dependent solubilisation and ligand binding affinity at CGRP and PTH1 receptors. Shown are the polymer concentrations used, solubilisation efficiency, and ligand binding affinity (pKd) under different copolymer environments. Data are mean ± s.e.m of n = 3 experiments performed in triplicate. In each case the pKd values in the nanodiscs were not significantly different to the pKd value in membranes when analysed by One-Way ANOVA with post-hoc Dunnett’s tests.

| **NLuc-PTH<sub>1</sub> receptor** |  |  |  |
| --- | --- | --- | --- |
| **Environment** | **Polymer concentration (w/v)** | **Solubilisation efficiency (%)** | **pK<sub>d</sub>** |
| **Membrane** | N/A | N/A | 7.30 ± 0.09 |
| **SMA2000** | 2.50% | 97.0 ± 1.5 | 7.54 ± 0.13 |
| **DIBMA-12** | 5.00% | 98.0 ± 1.1 | 6.97 ± 0.10 |
| **Sulfo-DIBMA** | 5.00% | 34.0 ± 10.2 | 7.06 ± 0.10 |
| **NLuc-CGRP receptor** |  |  |  |
| **Environment** | **Polymer concentration (w/v)** | **Solubilisation efficiency (%)** | **pK<sub>d</sub>** |
| **Membrane** | N/A | N/A | 9.32 ± 0.03 |
| **SMA2000** | 2.50% | 77.9 ± 5.9 | 9.74 ± 0.33 |
| **DIBMA-12** | 5.00% | 85.9 ± 2.1 | 9.50 ± 0.05 |
| **Sulfo-DIBMA** | 5.00% | 49.2 ± 3.5 | 9.32 ± 0.09 |

Although ligand affinity was retained, the copolymer environment had a marked impact on assay background. SMA 2000 in particular produced high non-specific binding for BODIPY-CGRP, consistent with strong peptide–polymer interactions driven by its smaller nanodiscs size and high anionic charge density. DIBMA-12 also showed elevated non-specific binding, but to a lesser extent than SMA 2000. In contrast, the electroneutral sulfo-DIBMALP exhibited minimal non-specific binding and closely resembled the membrane control. These observations indicate that, while all three copolymers preserve ligand affinity, polymer charge and chemistry can influence non-specific tracer binding at the CGRP receptor and therefore assay background.

### 3.3. DIBMA and sulfo-DIBMA encapsulated NLuc-CGRPR preserve competitive antagonist binding

Having confirmed that NLuc-CGRPR retained binding of BODIPY-CGRP in native-like nanodiscs, this enabled competitive ligand binding assays to be utilised using BODIPY-CGRP as tracer. SMALP was excluded from antagonist competition studies due to its high non-specific binding. Affinity purified NLuc-CGRPR in DIBMALP or sulfo-DIBMALP, together with NLuc-CGRPR in membranes as a reference, were subjected to equilibrium competition binding in the presence of 1 nM BODIPY-CGRP as tracer. Competing ligands tested covered three mechanistic classes: the truncated peptide antagonist CGRP_8-37_ (10 pM–10 µM), the small-molecule CLR-core antagonists olcegepant and rimegepant (100 fM–100 nM), and the extracellular domain (ECD)-targeting monoclonal antibody erenumab (1 pM–1 µM).

All four antagonists produced clear, concentration-dependent displacement of BODIPY-CGRP in membrane, DIBMALP and sulfo-DIBMALP preparations, demonstrating preserved antagonist binding in both nanodisc environments (shown in Figure 3). For olcegepant, rimegepant and erenumab, pKi values in DIBMALP and sulfo-DIBMALP were statistically indistinguishable from membrane values (pK_i_ values listed in Table 3), indicating that these nanodiscs maintained antagonist pharmacology across small-molecule and antibody classes. The only consistent deviation from membrane-derived values was observed for CGRP_8-37_ in both DIBMALP and sulfo-DIBMALP, suggesting a modest copolymer-dependent effect on this truncated peptide antagonist. Curiously, the affinity for both CGRP and CGRP_8-37_ in membranes were well over an order of magnitude less than reported for radioligand binding and indeed the direct binding of CGRP in the NanoBRET assay reported above. An assay-specific observation was that NLuc-CGRPR-DIBMALP retained a residual BRET signal at saturating antagonist concentrations, consistent with a weak, localised interaction between NLuc-CGRPR and BODIPY-CGRP that persisted after apparent displacement. Despite this residual signal, competition curves were well fitted and the derived pKi values did not significantly differ from membrane controls (as listed in Table 3), indicating preserved antagonist affinity.

**Figure 3.**
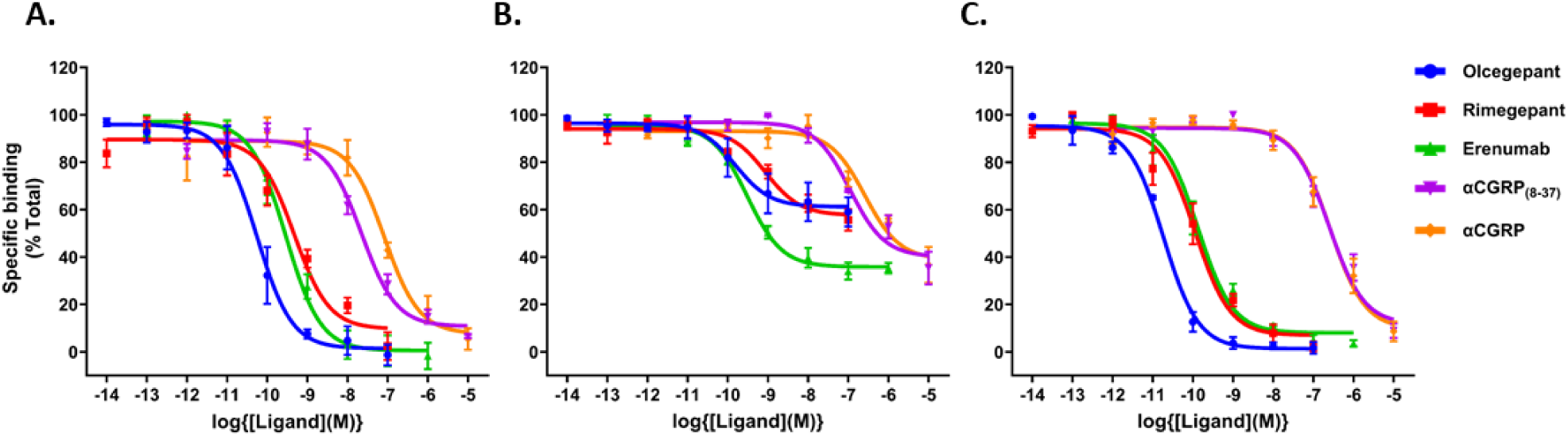
Equilibrium competition binding at membrane-bound, DIBMA- and sulfo-DIBMA-encapsulated NLuc-CGRPR. NanoBRET-based equilibrium competition binding performed using a fixed concentration of BODIPY-CGRP (1 nM) at (A) membrane-bound NLuc-CGRPR, (B) NLuc-CGRPR-DIBMALP, and (C) NLuc-CGRPR-sulfo-DIBMALP. Each panel shows displacement of BODIPY-CGRP by all four antagonists: CGRP, CGRP₈–₃₇ (10 pM–10 µM), Olcegepant, Rimegepant (100 fM–100 nM), and Erenumab (1 pM–1 µM). Curves represent mean competition binding profiles from three independent experiments performed in technical triplicate (n = 3).

**Table 3.** pKᵢ values derived from NanoBRET competition binding assays for antagonist ligands at NLuc-CGRPR in membrane, DIBMALP, and sulfo-DIBMALP. Data are presented as mean ± s.e.m. from n = 3 independent experiments performed in triplicate. pKi values were compared across each row and analysed by one-way ANOVA with Dunnett’s post-hoc tests, using membrane as the control condition. Statistical significance was accepted at P < 0.05 and is indicated by * where applicable.

| **Ligand** | **Membrane** | **DIBMALP** | **Sulfo-DIBMALP** |
| --- | --- | --- | --- |
| **Olcegepant** | 10.60 ± 0.19 | 10.45 ± 0.54 | 11.13 ± 0.04 |
| **Rimegepant** | 9.77 ± 0.17 | 9.59 ± 0.13 | 10.37 ± 0.26 |
| **Erenumab** | 10.03 ± 0.09 | 10.05 ± 0.12 | 10.34 ± 0.14 |
| **CGRP<sub>8-37</sub>** | 8.12 ± 0.09 | 7.45 ± 0.12* | 7.03 ± 0.14* |
| **αCGRP** | 7.55 ± 0.16 | 7.01 ± 0.3 | 7.03 ± 0.12 |

In summary, equilibrium competition binding showed that DIBMA- and sulfo-DIBMA-encapsulated NLuc-CGRPR retained antagonist pharmacology across peptide, small-molecule and antibody classes. Competition curves were well fitted and derived pKi values were comparable to membrane controls, indicating preserved antagonist competition for receptor binding. Although, copolymer chemistry potentially altered absolute BRET amplitudes in NLuc-CGRPR-DIBMALP conditions, these changes affected assay background and baseline signal but did not shift affinity estimates.

### 3.4. Sulfo-DIBMALP maintains G-protein coupling at CGRP receptor

To determine whether DIBMALP and sulfo-DIBMALP support an active intracellular receptor conformation, we assessed transducer engagement using Venus-tagged mini-Gα_s_ to measure direct G-protein association at CGRP receptor. This approach investigated whether the intracellular (C-terminal) NLuc tagged CGRP receptor (CGRPR-NLuc) remained competent to recruit G-protein in DIBMALP and sulfo-DIBMALP. G-protein coupling, using mini-Gαs-protein, was therefore examined in both DIBMALP and sulfo-DIBMALP preparations.

Canonical Gα_s_-protein engagement at CGRPR-NLuc was assessed using Venus-tagged mini-Gα_s_ in response to three CGRP-active peptide agonists: αCGRP, AM, and AM2. As shown in Figure 4, mini-Gα_s_ recruitment was clearly observed for CGRPR-NLuc encapsulated in sulfo-DIBMALP, whereas no detectable coupling occurred in DIBMALP, where the BRET signal declined towards baseline at higher ligand concentrations. h In contrast, sulfo-DIBMALP supported robust ligand-dependent recruitment of G-protein, with G-protein binding potencies (pEC₅₀) not significantly different from membrane-bound CGRPR-NLuc (Table 4), indicating that sulfo-DIBMALP preserves activation-competent receptor states.

**Figure 4.**
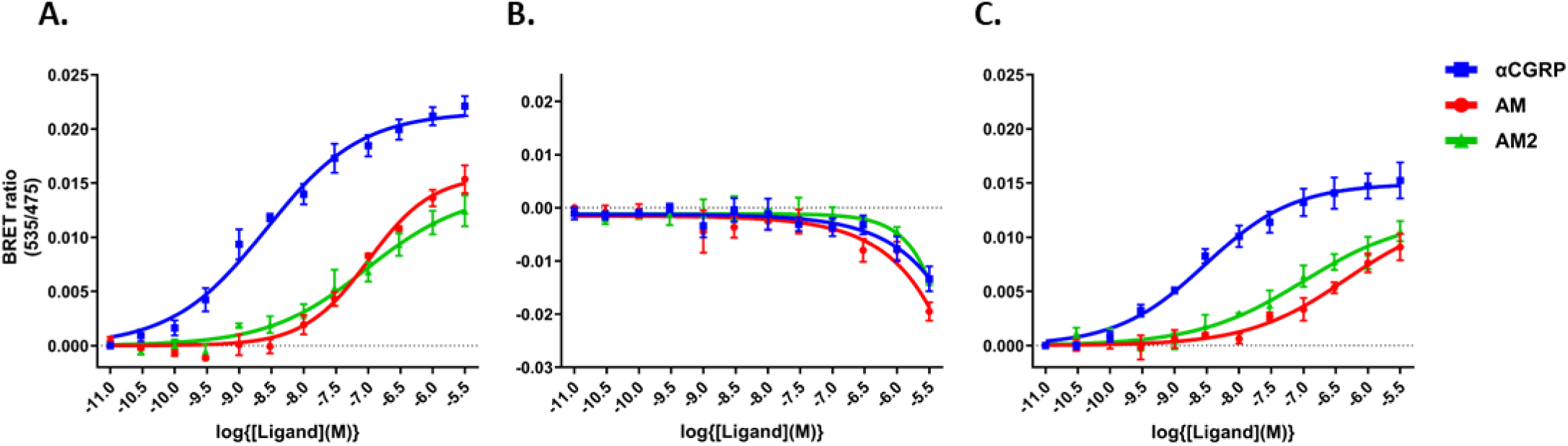
Mini-Gαs recruitment to CGRPR-NLuc is observed in sulfo-DIBMALP but not in DIBMALP. Equilibrium coupling between Venus–mini-Gαs and CGRPR-NLuc was assessed using NanoBRET assays in (A) membranes, (B) DIBMALP, and (C) sulfo-DIBMALP following stimulation with αCGRP, AM, or AM2. Data are presented as mean ± s.e.m. from three independent experiments (n = 3).

**Table 4.**
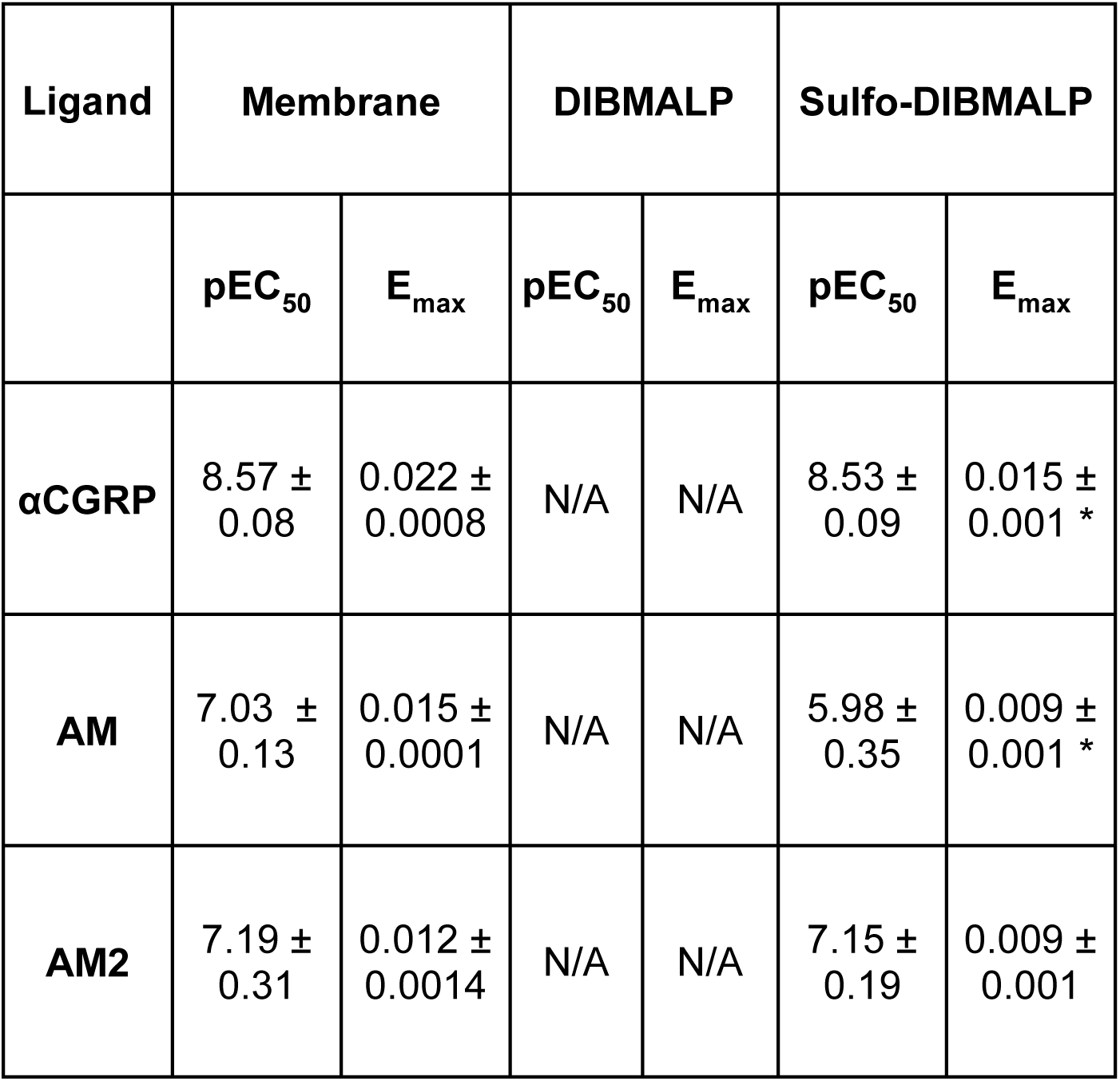
pEC₅₀ and Emax values for Venus–mini-Gαs coupling to CGRPR-NLuc following stimulation with αCGRP, adrenomedullin, or adrenomedullin-2/intermedin in membranes and sulfo-DIBMALP. Data are presented as mean ± s.e.m. from n = 3 independent experiments performed in triplicate. pEC₅₀ values were compared across each row and analysed by one-way ANOVA with Dunnett’s post-hoc tests, using membranes as the control condition. Statistical significance was accepted at P < 0.05 and is indicated by * where applicable.

### 3.5. Sulfo-DIBMALP maintains G-protein-dependent agonist affinity modulation at PTH_1_ receptor

After confirming that sulfo-DIBMA encapsulated CGRPR-NLuc can directly recruit Venus-mini-Gα_s_, we assessed whether G-protein binding allosterically modulated agonist affinity, using sulfo-DIBMALP-NLuc-PTH_1_R.

Two G-protein subtypes, mini-Gα_s_ and mini-Gα_q_, were examined because PTH_1_R can couple to both, and previous studies indicated subtype-specific differences in the active states they stabilise (38,39). As shown in Figure 5A, equilibrium saturation assays revealed that mini-Gα_s_ significantly increased BODIPY-PTH_1-34_ affinity by approximately six-fold compared with buffer alone, whereas mini-Gα_q_ did not generate a significant change. Competition binding assays (Figure 5B) showed a similar trend, with PTH_1-34_ affinity (pK_i_) significantly increasing ∼8.9-fold in the presence of mini-Gα_s_ and modestly increasing (∼1.7-fold) with mini Gα_q_, relative to buffer (Table 5).

**Figure 5.**
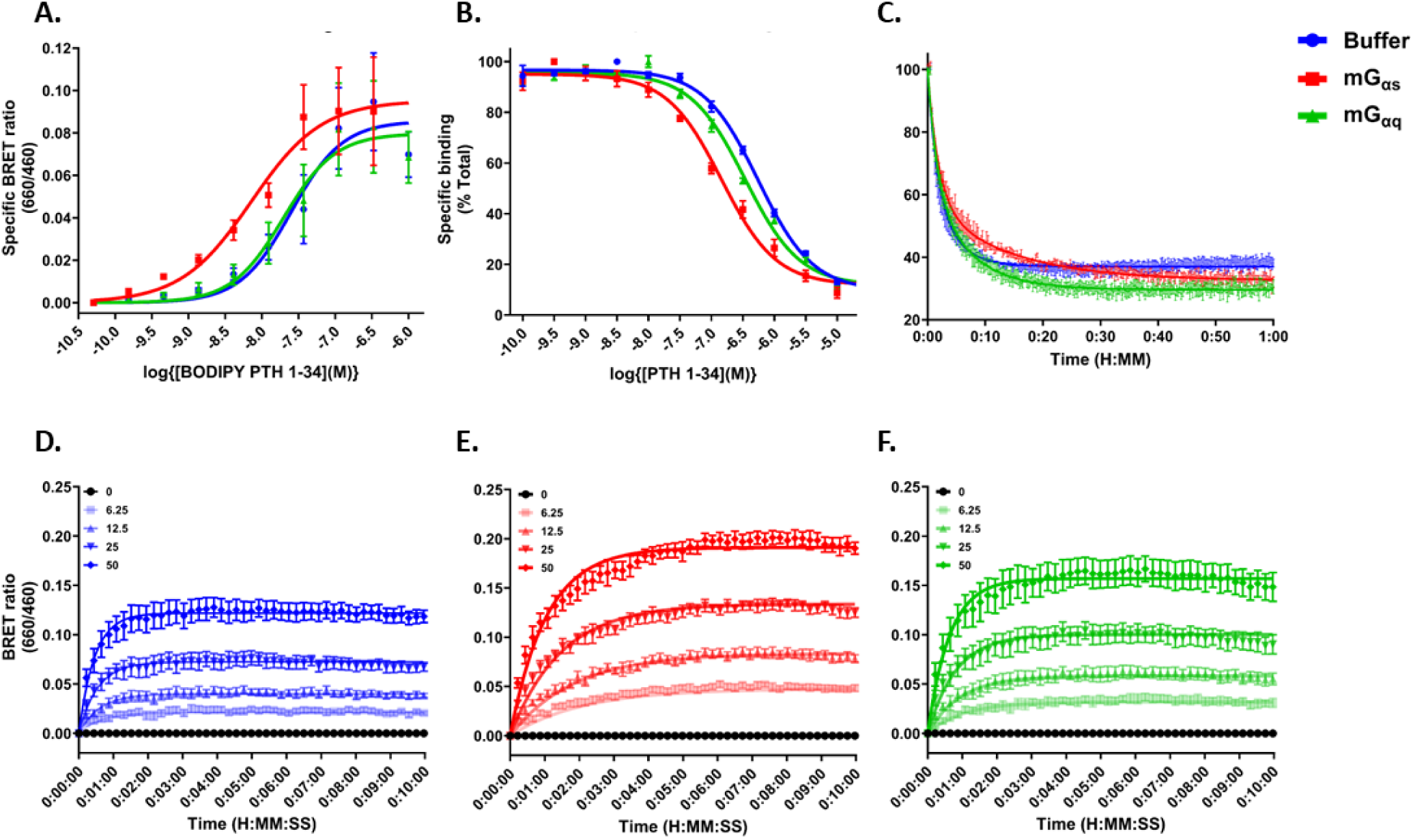
Equilibrium and kinetic binding of BODIPY-PTH_1-34_ at sulfo-DIBMALP-NLuc-PTH_1_R in the presence of mini-Gαs or mini-Gαq. NanoBRET-based equilibrium and kinetic binding analyses at sulfo-DIBMALP NLuc-PTH_1_R in the presence of mini-Gα proteins. (A) Equilibrium saturation binding of BODIPY-PTH_1-34_ in buffer (blue), mini-Gαs (red), or mini-Gαq (green), with specific binding defined by 10 μM DS69910557. (B) Equilibrium competition binding of unlabelled PTH_1-34_. (C) Dissociation of BODIPY-PTH_1-34_ initiated by 4 μM unlabelled PTH_1-34_. Kinetic association of BODIPY-PTH_1-34_ in the presence of (D) buffer, (E) mini-Gαs, and (F) mini-Gαq. Data are presented as mean ± s.e.m. from experiments performed in singlet (kinetic association), duplicate (kinetic dissociation), or triplicate for equilibrium saturation and competition binding.

**Table 5:** Binding constants from both kinetic and equilibrium binding assays performed at sulfo-DIBMALP encapsulated NLuc-PTH1R. Results are the mean ± s.e.m of experiments performed in singlet (Kinetic association binding), duplicate (Kinetic dissociation binding), or triplicates (Equilibrium saturation or competition binding). pK_d_ values were compared down a column. pK_d_ or pK_i_ values were analysed by One-Way ANOVA with post-hoc Dunnett’s tests, comparing buffer with mGαs and mGαq. Statistical significance was accepted when P < 0.05 and represented by an *.

| Kinetic Binding |  |  |  |  |  |  |  |  |  |  | Equilibrium Binding |  |
| --- | --- | --- | --- | --- | --- | --- | --- | --- | --- | --- | --- | --- |
|  | Association |  |  |  | Dissociation |  |  |  |  |  | Saturation | Competition |
|  | pK<sub>d</sub> | K<sub>on</sub><br>(10<sup>6</sup>M<sup>-1</sup>Min<sup>-1</sup>) | K<sub>off</sub><br>(Min<sup>-1</sup>) | RT<br>(minute) | pK<sub>d</sub><br>slow | pK<sub>d</sub><br>fast | K<sub>off</sub><br>slow | K<sub>off</sub><br>fast | RT<br>slow | RT<br>fast | pK<sub>d</sub> | pK<sub>i</sub> |
| Buffer | 6.95 ± 0.06 | 13.67 ± 2.94 | 1.53 ± 0.3 | 0.82 ± 0.17 | 7.61 ± 0.10 | 7.18 ± 0.10 | 0.29 ± 0.059 | 0.79 ± 0.029 | 3.67 ± 0.69 | 1.27 ± 0.05 | 7.58 ± 0.06 | 6.40 ± 0.03 |
| mGα<sub>s</sub> | 7.42 ± 0.05 * | 11.36 ± 1.50 | 0.43 ± 0.06 | 2.55 ± 0.37 | 8.18 ± 0.07 * | 7.31 ± 0.07 | 0.07 ± 0.003 | 0.54 ± 0.052 | 14.0 ± 0.68 | 1.90 ± 0.20 | 8.36 ± 0.05 * | 7.35 ± 0.05 * |
| mGα<sub>q</sub> | 7.20 ± 0.05 * | 15.61 ± 3.84 | 1.07 ± 0.36 | 1.53 ± 0.53 | 7.90 ± 0.13 | 7.32 ± 0.13 | 0.17 ± 0.023 | 0.64 ± 0.030 | 6.2 ± 0.74 | 1.57 ± 0.07 | 7.69 ± 0.04 | 6.64 ± 0.04 * |

To further characterise these effects, kinetic binding assays were performed to determine how mini-Gα proteins modulate ligand binding for receptors encapsulated within sulfo-DIBMALP. As shown in Figure 5C, dissociation data fitted a two-phase exponential model, consistent with two receptor populations reported previously (40–43). Mini-Gα_s_ had little effect on the fast phase (k_fast_) but significantly slowed the slow phase (k_slow_), increasing ligand residence time by ∼3.8-fold, while mini-Gα_q_ produced a smaller ∼1.7-fold increase (Table 5). Kinetic association data Figure 5D, 5E and 5F, best described by a single-phase model, reinforced these differences, showing the greatest affinity increase with mini-Gα_s_ over mini-Gα_q_. This increase arises primarily from reduced ligand dissociation (k_off_), rather than faster association (k_on_). Calculated pK_d_ values derived from k_on_/k_off_ correlated well with equilibrium saturation data (Table 5). Together, these data demonstrate that sulfo-DIBMALP supports high-affinity ligand-binding and enables G-protein-dependent affinity modulation for PTH_1_ receptor.

## 4. DISCUSSION

Two uncertainties have shaped our rationale for investigating amphipathic copolymers in functional GPCR studies. First, whether the charged hydrophilic copolymer subunit may introduce non-specific interactions that compromise quantitative pharmacology. Second whether copolymer nanodiscs can preserve the intracellular conformational dynamics required for G-protein recruitment and G-protein-dependent allosteric modulation of agonist binding affinity. These considerations are particularly important for class B1 GPCRs, whose activation involves lipid-mediated allosteric transitions and extensive intracellular rearrangements (44,45). Our study addresses these points by defining a mechanistic framework that links polymer chemistry to GPCR signalling competence.

By comparing SMA 2000, DIBMA and sulfo-DIBMA, this study systematically dissected the contributions of aromaticity, charge density and membrane perturbation to receptor behaviour. The copolymer panel in this study investigated parameters most likely to influence class B1 GPCR stability, peptide ligand binding ability and intracellular signalling competence.

Across all three copolymers, NanoBRET ligand binding assays showed that high-affinity peptide binding was preserved, with pK_d_ values comparable to those in native membranes (Table 2). This confirms that SMA 2000, DIBMA and sulfo-DIBMA all maintain the extracellular architecture and orthosteric ligand binding pocket of PTH_1_ and CGRP receptors. These finding extend previous observations that aliphatic copolymers support ligand binding and stabilise active-state transitions in other GPCRs (11, 44–46). However, the requirement to remove free polymer prior to binding assays reinforces the susceptibility of peptide ligands to non-specific interactions with maleic acid-containing copolymers, consistent with earlier reports (49,50).

At physiological pH, CGRP carries a net positive charge (approximately +3) and PTH_1-34_ is also cationic (approximately +1). Peptide net charges are estimated from their primary sequences using standard amino-acid pK_a_ values (51,52). In contrast, SMA 2000 and DIBMA are strongly anionic due to the full deprotonation of maleic acid moieties (22,23). This generates a pronounced electrostatic asymmetry that drives non-specific peptide-polymer association, where CGRP and PTH_1-34_ with their positive charge are more prone to sequestration by these polymers. This consequently reduces the free available peptide ligand concentration and potentially distorts apparent potency and kinetics. In contrast to SMA and DIBMA, sulfo-DIBMA is not an anionic copolymer but a zwitterionic one. Each monomer contains a permanently charged quaternary ammonium group balanced by a sulfonate, giving the polymer a net zero charge at all pH values (25,53). This internal charge balance, together with hydration shell characteristic of sulfobetanine-type zwitterions (53), generates an effectively electroneutral polymer surface that lacks the electrostatic hotspots responsible for CGRP or PTH_1-34_ tracers associating with the polymer belt in the sulfo-DIBMALP (Figure 2). This physiochemical profile accounts for the observed minimal non-specific peptide association, where the non-specific binding values of sulfo-DIBMALP are comparable to those of the membrane only control. This demonstrated that sulfo-DIBMA does not engage in electrostatic peptide interactions and therefore preserves ligand-receptor pharmacology without polymer-mediated artefacts.

Despite all copolymers supporting ligand binding, with polymer-dependent differences in non-specific background, the NanoBRET G-protein recruitment assays revealed a functional divergence. Only sulfo-DIBMA preserved intracellular signalling competence, supporting both direct G-protein recruitment and G-protein dependent allosteric modulation of agonist binding. This distinction exposes a fundamental principle: the polymer properties required to maintain extracellular ligand recognition are not the same as those required to preserve intracellular conformational dynamic required for G-protein coupling. This interpretation aligns with the inability of DIBMA to support G-protein coupling for the ghrelin receptor (14) but contradicts with the report of SMALP-supported ghrelin receptor coupling to G-protein (14). This difference potentially reflects methodological differences, as our approach extracted human class B1 receptors directly from mammalian cell membranes not from detergent-treated proteoliposomes, avoiding any detergent-induced conformational bias.

The superior performance of sulfo-DIBMA identifies charge neutrality as critical determinants of class B1 GPCR signalling competence. Substituting the maleic acid group with an electroneutral maleimide derivative (25), while keeping DIBMA’s fluidity-preserving aliphatic backbone (54) that maintain receptor flexibility (48), allows sulfo-DIBMA to maintain the intracellular conformational freedom needed for G-protein recruitment and allosteric modulation. This mechanistic insight explains why ligand binding is broadly preserved across polymers, yet G-protein recruitment competence is restricted to sulfo-DIBMALP. It also provides a conceptual basis for selecting copolymer chemistries tailored to the signalling requirements of different GPCR families.

Despite these advances, some important questions remain. It was beyond the scope of the study to determine whether different polymers bias the lipid species co-extracted with the receptor. Although lipidomic analyses suggest that polymer nanodiscs broadly reflect the native membrane environment (55,56), polymer-dependent lipid selectivity has not been systematically examined. In addition, lipid-swapping has been observed for lipid-only discs (57) and for rhomboid proteases (58) but has not been demonstrated for GPCR-lipid nanodiscs. Given the established role of specific lipids in stabilising class B1 GPCR activation (59), future work should test whether exogenous lipids can be introduced to modulate receptor function. Extending this approach to other GPCR effectors, such as G-protein-coupled receptor kinases (60) and β-arrestin (61), whose membrane interactions are lipid-dependent (62), will be essential for determining whether copolymer nanodiscs can support the full complement of family B GPCR signalling pathways. Disc size may also be a factor in stabilising different protein conformations (48,63). Our data show other factors also need to be considered as sulfo-DIBMA gives discs the same diameter as SMA and which are only half the size of those with DIBMA-12 (Table 1).

We observed that in competition experiments our BODIPY-CGRP behaved anomalously with respect to the peptide displacing ligands. It may be that the BODIPY group is able to interact with the TM regions of the CGRP receptor, although it behaves as expected in direct binding and cAMP stimulation (Supplementary Figure 1).

In summary, our data suggest that copolymer chemistry can influence the signalling competence of purified class B1 GPCRs. While SMA 2000 and DIBMA-12 preserve ligand binding, only sulfo-DIBMA appears to maintain the intracellular conformational freedom required for G-protein recruitment and G-protein dependent allostery. Taken together, these findings indicate that sulfo-DIBMA offers a particularly compatible platform for studying class B1 GPCRs in native-like environments and provides a potential mechanistic framework to guide rational copolymer selection in future pharmacological, biophysical and structural investigations.

## Abbreviations

AM: adrenomedullin
AM2: adrenomedullin 2 (intermedin)
BODIPY: boron-dipyrromethene
BRET: bioluminescence resonance energy transfer
cAMP: cyclic adenosine monophosphate
cDNA: complementary DNA
CGRP: calcitonin gene-related peptide
CGRPR: calcitonin gene-related peptide receptor
CLR: calcitonin receptor-like receptor
CTR: calcitonin receptor
DIBMA: poly(diisobutylene-*alt*-maleic acid)
DIBMALP: DIBMA lipid particles
DMEM: Dulbecco’s Modified Eagle Medium
DMEM/F12: Dulbecco’s Modified Eagle Medium/Nutrient Mixture F-12
DTT: dithiothreitol
*E. coli*: Escherichia coli
EDTA: ethylenediaminetetraacetic acid
EGTA: ethylene glycol-bis(β-aminoethyl ether)-N,N,N′,N′-tetraacetic acid
FBS: foetal bovine serum
G418: geneticin
GDP: guanosine diphosphate
GLP1R: glucagon-like peptide-1 receptor
GNTI-: N-acetylglucosaminyltransferase I deficient
GPCRs: G-protein-coupled receptors
HA: hemagglutinin
HEK 293S: human embryonic kidney 293S cells
HEPES: 4-(2-hydroxyethyl)-1-piperazineethanesulfonic acid
His_10_: deca-histidine
IMP: Integral membrane protein
IPTG: isopropyl β-D-1-thiogalactopyranoside
LVF: Linear Variable Filter
Mini-Gα_q_: minimal G-protein α subunit (Gq family)
mini-Gα_s_: minimal G-protein α subunit (Gs family)
NanoBRET: nano-bioluminescence resonance energy transfer
Ni-NTA: nickel nitrilotriacetic acid
NLuc: NanoLuciferase
OD_600_: optical density at 600 nm
P2A: porcine teschovirus-1 2A element
PEI: polyethylenimine
Pra: propargylglycine
PTH1: parathyroid hormone 1
PTH_1-34_: parathyroid hormone fragment 1–34
PTH_1_R: parathyroid hormone 1 receptor
RAMP1: receptor activity modifying protein 1
RPM: revolutions per minute
RT: room temperature
SEC: size-exclusion chromatography
SDS: sodium dodecyl sulfate
SDS-PAGE SDS: polyacrylamide gel electrophoresis
SMA: poly(styrene-*co*-maleic acid)
SMALP: SMA lipid particle
Sulfo-DIBMA: sulfobetaine- poly(diisobutylene-*alt*-maleic acid)
Sulfo-DIBMALP: sulfo-DIBMA lipid particles
TAMRA: tetramethylrhodamine
TetO: tetracycline operator
TetR: tetracycline repressor
TM6: transmembrane helix 6

## Data Availability statement

All data reported in this paper will be shared by the corresponding author upon reasonable request.

## Acknowledgements

JG was supported by an MIBTP CASE award with Inocardia. PK was supported by a BBSRC Discovery fellowship (BB/W00934X/1).

## COI/Notes

The authors declare no competing financial interest

