## Supplementary Figures for "Sulfo-DIBMA encapsulation uniquely preserves signalling-competent active states of Class B1 GPCRs, CGRP and PTH1 receptors, in native-like nanodiscs"

αCGRP: ACDTATCVTHRLAGLLSRSGGVVKNNFVPTNVGSKAF
Fluorescent αCGRP: AC[Pra(BODIPY 630/650)]TATCVTHRLAGLLSRSGGVVKNNFVPTNVGSKAF

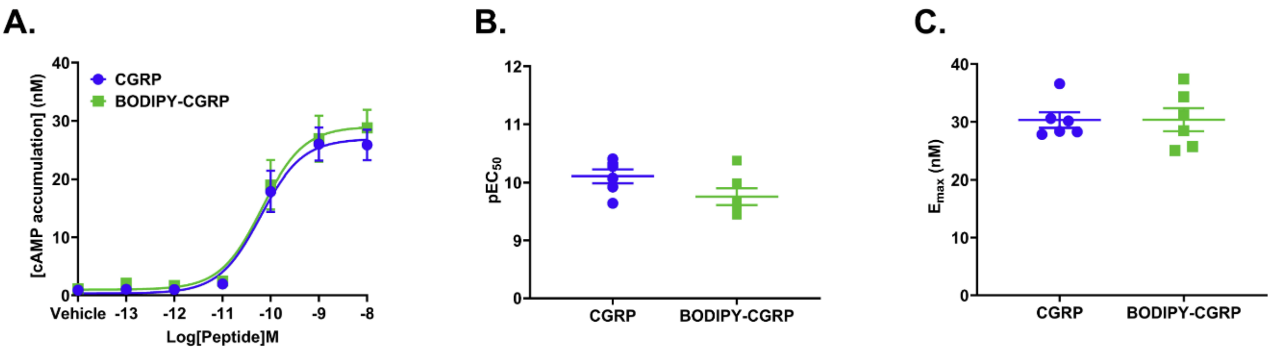


**Supplementary Figure 1. Fluorescently derivatised α-CGRP with BODIPY 630/650 and functional pharmacological characterisation of BODIPY-CGRP in NLuc-CGRPR-expressing cells.**

Amino acid sequence of wild-type αCGRP and BODIPY-derivatised, fluorescently labelled αCGRP (BODIPY-CGRP). The derivatised amino acid is highlighted in green. Pra=L-propargylglycine. cAMP accumulation as second messenger assay performed to determine potency of BODIPY-CGRP with WT-CGRP receptor in stably bi-gene NLuc-CGRPR expressing HEK 293S cells. Cells stably expressing NLuc-CGRPR were exposed to induction medium containing tetracycline (2 µg/mL). (A) Concentration–response curves for α-CGRP (100 fM–10 nM) and BODIPY-CGRP (100 fM–10 nM). (B) $pEC_{50}$ values for α-CGRP and BODIPY-CGRP. (C) $E_{max}$ values for α-CGRP and BODIPY-CGRP. All values were calculated from individual experiments, and error bars represent the mean ± s.e.m. Statistical significance was determined relative to control using an unpaired t-test ($p < 0.05$).

$PTH_{1-34}$: SVSEIQLMHNLGKHLNSMERVEWLRKKLQDVHNF
BODIPY-$PTH_{1-34}$: SVSEIQLMHNLG[Lys(BODIPY 630/650)]HLNSMERVEWLRKKLQDVHNF

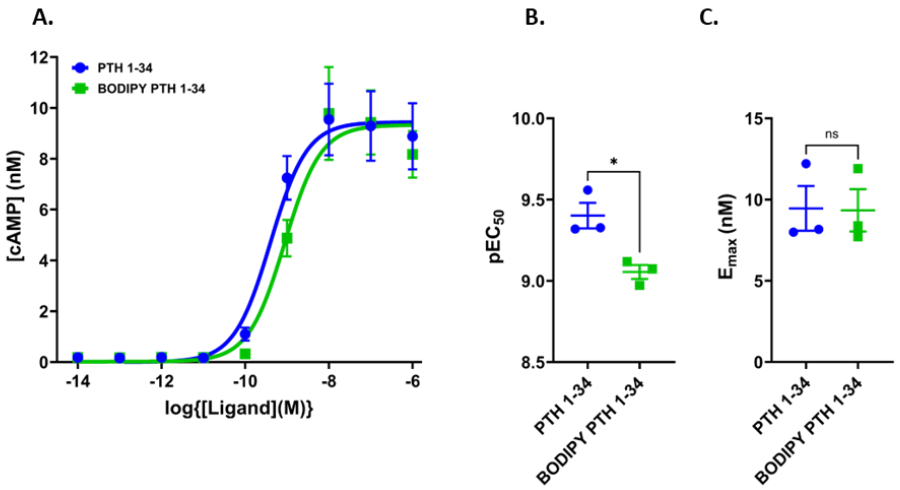


**Supplementary Figure 2. Fluorescently tagged $PTH_{1-34}$ with BODIPY 630/650 and functional pharmacological characterisation of BODIPY-$PTH_{1-34}$ in NLuc-PTH1R-expressing cells.** Amino acid sequence of wild-type $PTH_{1-34}$ and $Lys^{13}$ derivatised, fluorescently (BODIPY) labelled $PTH_{1-34}$ (BODIPY-$PTH_{1-34}$). The derivatised amino acid is highlighted in green. cAMP accumulation as second messenger assay performed to determine potency of BODIPY-$PTH_{1-34}$ with WT-$PTH_{1-34}$ in stably NLuc-$PTH_1$R expressing HEK 293S cells. Cells stably expressing NLuc-$PTH_1$R were exposed to induction medium containing tetracycline (2 µg/mL). (A) Concentration–response curves for $PTH_{1-34}$ (100 fM–10 nM) and BODIPY-$PTH_{1-34}$ (100 fM–10 nM). (B) $pEC_{50}$ values for $PTH_{1-34}$ and BODIPY-$PTH_{1-34}$. (C) $E_{max}$ values for $PTH_{1-34}$ and BODIPY-$PTH_{1-34}$. All values were calculated from individual experiments, and error bars represent the mean ± s.e.m. Statistical significance was determined relative to control using an unpaired t-test ($p < 0.05$).

### CGRP receptor

START- HA signal peptide- His$_{10}$ TAG- TEV- HA TAG- CLR- 15 A.A linker- AVI TAG- GSG- 2PA- HA signal peptide-FLAG TAG- RAMP1- STOP

MKTIIALSYIFCLVFAHHHHHHHHHHENLYFQSYPYDVPDYAAELEESPEDSIQLGVTRNKIMTAQYEC
YQKIMQDPIQQAEGVYCNRTWDGWLCWNDVAAGTESMQLCPDYFQDFDPSEKVTKICDQDGNWF
RHPASNRTWTNYTQCNVNTHEKVKTALNLFYLTIIGHGLSIASLLISLGIFFYFKSLSCQRITLHKNLFFS
FVCNSVVTIIHLTAVANNQALVATNPVSCKVSQFIHLYLMGCNYFWMLCEGIYLHTLIVVAVFAEKQHL
MWYYFLGWGFPLIPACIHAIARSLYYNDNCWISSDTHLLYIIHGPICAALLVNLFFLLNIVRVLITKLKVTH
QAESNLYMKAVRATLILVPLLGIEFVLIPWRPEGKIAEEVYDYIMHILMHFQGLLVSTIFCFFNGEVQAIL
RRNWNQYKIQFGNSFSNSEALRSASYTVSTISDGPGYSHDCPSEHLNGKSIHDIENVLLKPENLYNG
GGGSGGGGSGGGGSGLNDIFEAQKIEWHEGSGATNFSLLKQAGDVEENPGPKTIIALSYIFCLVFAD
YKDDDDKCQEANYGALLRELCLTQFQVDMEAVGETLWCDWGRTIRSYRELADCTWHMAEKLGCFW
PNAEVDRFFLAVHGRYFRSCPISGRAVRDPPGSILYPFIVVPITVTLLVTALVVWQSKRTEGIV-

### NLuc-CGRPR

START- HA signal peptide- His$_{10}$ TAG- TEV- HA TAG- NLuc- GSG- CLR- 15 A.A linker- Twin Strep II tag- GSG- P2A- HA signal peptide- FLAG TAG- RAMP1- STOP

MKTIIALSYIFCLVFAHHHHHHHHHHENLYFQSYPYDVPDYAMVFTLEDFVGDWRQTAGYNLDQVLE
QGGVSSLFQNLGVSVTPIQRIVLSGENGLKIDIHVIIPYEGLSGDQMGQIEKIFKVVYPVDDHHFKVILH
YGTLVIDGVTPNMIDYFGRPYEGIAVFDGKKITVTGTLWNGNKIIDERLINPDGSLLFRVTINGVTGWRL
CERILAGSGAELEESPEDSIQLGVTRNKIMTAQYECYQKIMQDPIQQAEGVYCNRTWDGWLCWNDV
AAGTESMQLCPDYFQDFDPSEKVTKICDQDGNWFRHPASNRTWTNYTQCNVNTHEKVKTALNLFYL
TIIGHGLSIASLLISLGIFFYFKSLSCQRITLHKNLFFSFVCNSVVTIIHLTAVANNQALVATNPVSCKVSQ
FIHLYLMGCNYFWMLCEGIYLHTLIVVAVFAEKQHLMWYYFLGWGFPLIPACIHAIARSLYYNDNCWIS
SDTHLLYIIHGPICAALLVNLFFLLNIVRVLITKLKVTHQAESNLYMKAVRATLILVPLLGIEFVLIPWRPEG
KIAEEVYDYIMHILMHFQGLLVSTIFCFFNGEVQAILRRNWNQYKIQFGNSFSNSEALRSASYTVSTISD
GPGYSHDCPSEHLNGKSIHDIENVLLKPENLYNGGGGSGGGGSGGGGSSAWSHPQFEKGGGSGG
GSGGSAWSHPQFEKGSGATNFSLLKQAGDVEENPGPKTIIALSYIFCLVFADYKDDDDKCQEANYGA
LLRELCLTQFQVDMEAVGETLWCDWGRTIRSYRELADCTWHMAEKLGCFWPNAEVDRFFLAVHGR
YFRSCPISGRAVRDPPGSILYPFIVVPITVTLLVTALVVWQSKRTEGIV-

### CGRPR-NLuc

START- HA signal peptide- His$_{10}$ TAG- TEV- AVI TAG- CLR- GSG- NLuc- GSG- P2A- HA signal peptide- HA TAG- RAMP1- STOP

MKTIIALSYIFCLVFAHHHHHHHHHHENLYFQSGLNDIFEAQKIEWHEAELEESPEDSIQLGVTRNKIMT
AQYECYQKIMQDPIQQAEGVYCNRTWDGWLCWNDVAAGTESMQLCPDYFQDFDPSEKVTKICDQD
GNWFRHPASNRTWTNYTQCNVNTHEKVKTALNLFYLTIIGHGLSIASLLISLGIFFYFKSLSCQRITLHK
NLFFSFVCNSVVTIIHLTAVANNQALVATNPVSCKVSQFIHLYLMGCNYFWMLCEGIYLHTLIVVAVFAE
KQHLMWYYFLGWGFPLIPACIHAIARSLYYNDNCWISSDTHLLYIIHGPICAALLVNLFFLLNIVRVLITKL
KVTHQAESNLYMKAVRATLILVPLLGIEFVLIPWRPEGKIAEEVYDYIMHILMHFQGLLVSTIFCFFNGEV
QAILRRNWNQYKIQFGNSFSNSEALRSASYTVSTISDGPGYSHDCPSEHLNGKSIHDIENVLLKPENL
YNGSGMVFTLEDFVGDWRQTAGYNLDQVLEQGGVSSLFQNLGVSVTPIQRIVLSGENGLKIDIHVIIP
YEGLSGDQMGQIEKIFKVVYPVDDHHFKVILHYGTLVIDGVTPNMIDYFGRPYEGIAVFDGKKITVTGT
LWNGNKIIDERLINPDGSLLFRVTINGVTGWRLCERILAGSGATNFSLLKQAGDVEENPGPKTIIALSYI
FCLVFAYPYDVPDYACQEANYGALLRELCLTQFQVDMEAVGETLWCDWGRTIRSYRELADCTWHM
AEKLGCFWPNAEVDRFFLAVHGRYFRSCPISGRAVRDPPGSILYPFIVVPITVTLLVTALVVWQSKRTE
GIV-

**Supplementary Figure 3. CGRP receptor (CLR-P2A-RAMP1) construct designs and corresponding amino acid sequences used in this study.**

### $PTH_1$ receptor

START- HA signal peptide- His10 TAG- TEV- HA TAG- GSG- (26-593)PTH1R- 17 A.A linker- Twin Strep TAG- STOP

MKTIIALSYIFCLVFAHHHHHHHHHHENLYFQSYPYDVPDYAGSGVDADDVMTKEEQIFLLHRAQAQC
EKRLKEVLQRPASIMESDKGWTSASTSGKPRKDKASGKLYPESEEDKEAPTGSRYRGRPCLPEWDH
ILCWPLGAPGEVVAVPCPDYIYDFNHKGHAYRRCDRNGSWELVPGHNRTWANYSECVKFLTNETRE
REVFDRLGMIYTVGYSVSLASLTVAVLILAYFRRLHCTRNYIHMHLFLSFMLRAVSIFVKDAVLYSGATL
DEAERLTEEELRAIAQAPPPPATAAAGYAGCRVAVTFFLYFLATNYYWILVEGLYLHSLIFMAFFSEKK
YLWGFTVFGWGLPAVFVAVWVSVRATLANTGCWDLSSGNKKWIIQVPILASIVLNFILFINIVRVLATKL
RETNAGRCDTRQQYRKLLKSTLVLMPLFGVHYIVFMATPYTEVSGTLWQVQMHYEMLFNSFQGFFV
AIIYCFCNGEVQAEIKKSWSRWTLALDFKRKARSGSSSYSYGPMVSHTSVTNVGPRVGLGLPLSPRL
LPTATTNGHPQLPGHAKPGTPALETLETTPPAMAAPKDDGFLNGSCSGLDEEASGPERPPALLQEE
WETVMGGGGSGGGGSGGGGSSAWSHPQFEKGGGSGGGSGGSAWSHPQFEK-

### NLuc-$PTH_1R$

START- HA signal peptide- $His_{10}$ TAG- TEV- HA TAG- NLuc- GSG- (26-593)$PTH_1R$- 17 A.A linker- Twin Strep TAG- STOP

MKTIIALSYIFCLVFAHHHHHHHHHHENLYFQSYPYDVPDYAMVFTLEDFVGDWRQTAGYNLDQVLE
QGGVSSLFQNLGVSVTPIQRIVLSGENGLKIDIHVIIPYEGLSGDQMGQIEKIFKVVYPVDDHHFKVILH
YGTLVIDGVTPNMIDYFGRPYEGIAVFDGKKITVTGTLWNGNKIIDERLINPDGSLLFRVTINGVTGWRL
CERILAGSGVDADDVMTKEEQIFLLHRAQAQCEKRLKEVLQRPASIMESDKGWTSASTSGKPRKDKA
SGKLYPESEEDKEAPTGSRYRGRPCLPEWDHILCWPLGAPGEVVAVPCPDYIYDFNHKGHAYRRCD
RNGSWELVPGHNRTWANYSECVKFLTNETREREVFDRLGMIYTVGYSVSLASLTVAVLILAYFRRLH
CTRNYIHMHLFLSFMLRAVSIFVKDAVLYSGATLDEAERLTEEELRAIAQAPPPPATAAAGYAGCRVAV
TFFLYFLATNYYWILVEGLYLHSLIFMAFFSEKKYLWGFTVFGWGLPAVFVAVWVSVRATLANTGCW
DLSSGNKKWIIQVPILASIVLNFILFINIVRVLATKLRETNAGRCDTRQQYRKLLKSTLVLMPLFGVHYIVF
MATPYTEVSGTLWQVQMHYEMLFNSFQGFFVAIIYCFCNGEVQAEIKKSWSRWTLALDFKRKARSG
SSSYSYGPMVSHTSVTNVGPRVGLGLPLSPRLLPTATTNGHPQLPGHAKPGTPALETLETTPPAMAA
PKDDGFLNGSCSGLDEEASGPERPPALLQEEWETVMGGGGSGGGGSGGGGSSAWSHPQFEKGG
GSGGGSGGSAWSHPQFEK-

**Supplementary Figure 4. $PTH_1$ receptor construct designs and corresponding amino acid sequences used in this study.**

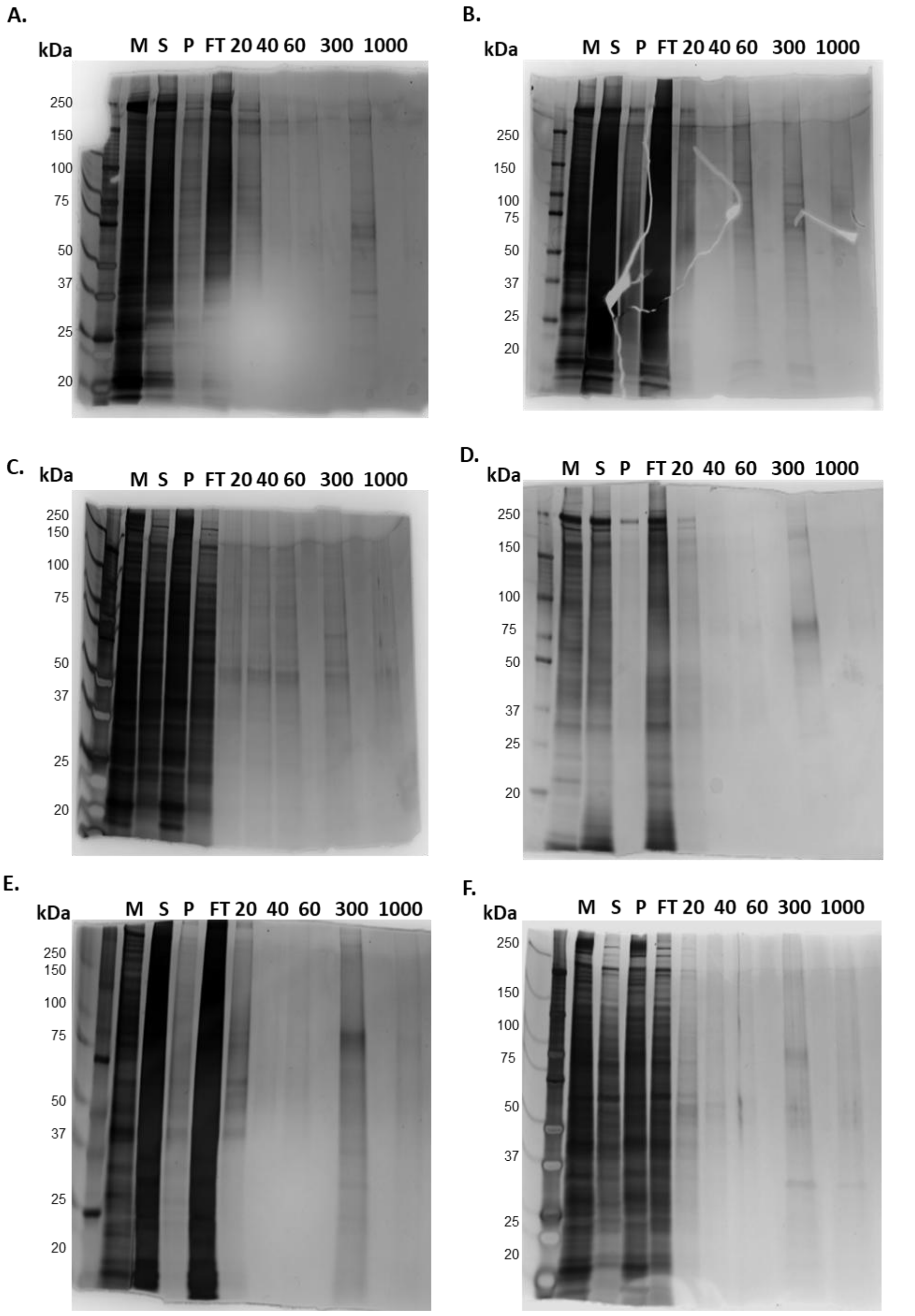


**Supplementary Figure 5. SDS-PAGE analysis of CGRP receptor and PTH$_1$ receptor solubilised and affinity-purified fractions in SMALP, DIBMALP and sulfo-DIBMALP.** Recombinant human CGRP and PTH$_1$ receptors stably overexpressed in HEK 293S GnTI$^-$ TetR cell membranes were used to assess solubilisation and affinity-purification profiles following extraction with 2.5% SMA 2000, 5% DIBMA, or 5% sulfo-DIBMA copolymers. Representative silver stained SDS PAGE gels are shown (n = 3). Panels: (A) CGRPR-SMALP; (B) CGRPR-DIBMALP; (C) CGRPR-sulfo-DIBMALP; (D) PTH$_1$R-SMALP; (E) PTH$_1$R-DIBMALP; (F) PTH$_1$R-sulfo-DIBMALP. Lane annotations: M = membrane; S = solubilised fraction; P = insoluble pellet; FT = affinity-purification flow-through. Imidazole concentrations (mM) for wash and elution steps are indicated (20, 40, 60, 300, 1000). Theoretical molecular weights: NLuc-PTH$_1$R ≈ 92 kDa; NLuc-CLR ≈ 74 kDa.
